# State-dependent differences in the frequency of TMS-evoked potentials between resting and active states

**DOI:** 10.1101/614826

**Authors:** Candice T. Stanfield, Martin Wiener

## Abstract

Previous evidence suggests different cortical areas naturally oscillate at distinct frequencies, reflecting tuning properties of each region. The concurrent use of transcranial magnetic stimulation (TMS) and electroencephalography (EEG) has been used to perturb cortical regions, resulting in an observed post-stimulation response that is maximal at the natural frequency of that region. However, little is known about the spatial extent of TMS-induced activation differences in cortical regions when comparing resting state (passive) versus active task performance. Here, we employed TMS-EEG to directly perturb three cortical areas in the right hemisphere while measuring the resultant changes in maximal evoked frequency in healthy human subjects during a resting state (N=12) and during an active sensorimotor task (N=12). Our results revealed that the brain engages a higher dominant frequency mode when actively engaged in a task, such that the frequency evoked during a task is consistently higher across cortical regions, regardless of the region stimulated. These findings suggest that a distinct characteristic of active performance versus resting state is a higher state of natural cortical frequencies.

## Introduction

The influence of task-evoked activation and behavior on the modification of spontaneously occurring patterns of neural activity remains a fundamental question in neuroscience. For decades, non-invasive brain stimulation techniques, such as transcranial magnetic stimulation (TMS), have been used to modulate neural activity in humans and other mammals. Furthermore, in numerous reports, concurrent TMS and electroencephalography (EEG) has been employed to examine cortical reactivity and connectivity. A variety of research using TMS, and some using concurrent TMS-EEG, demonstrates TMS-evoked behavioral and neural effects that are dependent on whether the subject is engaged in a task or not (Johnson, et al. 2012; Thut, et al. 2003; 2011; Romero, et al, 2019;Silvanto, et al. 2007;2008; Romei, et al. 2008; Massimini, et al. 2010;Miniussi, et al. 2010; Romei, et al. 2016; Petrichella, et al. 2017), as well as differences in neural activation during wakeful versus sleeping states (Massimini, et al. 2005).

In neural stimulation research, more is known about the influence exogenous factors have on the brain’s electrical response to TMS (frequency and intensity of stimulation; positioning and orientation of the stimulation coil), as opposed to endogenous factors (e.g., global brain state). However, over the past decade, there has been an emergence of research using concurrent TMS-EEG to investigate the influence of endogenous factors on neural response. One such study observed an increase in amplitude and spatial spread during the performance of a short-term memory task (Johnson, et al. 2012). Moreover, the observed task-related excitability increased as a result of stimulation to the cortical area, including spread of TMS-evoked currents to functionally connected areas. Globally, the dominant frequency recorded at the scalp matched that of the stimulated area. Yet, local cortical areas oscillated at a rate closer to its own natural frequency, even when not directly stimulated. Lower-frequency oscillatory peaks were observed in the frontal and parietal cortex (7 Hz in the theta band, and 10 Hz in the alpha band, respectively), reflecting synchronization of local cortical oscillations to parallel networks engaged in task performance. Similar results were reported in a study that provided the first direct evidence for causal entrainment of brain oscillations by short rhythmic TMS bursts while recording resultant EEG responses (Thut, et al. 2011). The TMS entrainment evoked spatially specific and frequency-specific oscillatory signatures that mimic naturally occurring task-related modulations that are of functional significance. Overall, these task-dependent changes exemplify the importance of further investigation into the influence of endogenous factors, such as global brain state.

Task-dependent changes have also been observed at the single-cell level, with concurrent single-pulse TMS administered to awake rhesus monkeys (Romero, et al. 2019). During performance of a visually-guided grasping task, action potentials in individual neurons within the parietal area PFG were recorded extracellularly, while either low intensity (60% of the resting Motor Threshold; rMT) or high intensity (120% of the rMT) stimulation was being administered. Unlike in previous observations of anesthetized animals, single-pulse stimulation induced a highly localized and transient excitation followed by reduced activity, corresponding with a significantly longer grasping time. Thus, the stimulation interfered with task-related activity in parietal neurons, while simultaneously causing behavioral effects. Additionally, the stimulation induced a highly localized and short-lived excitation of single neurons in the parietal cortex; however, the TMS-induced activity and task-related activity did not linearly summate in the PFG neurons. As such, the spatial spread of TMS-induced spiking activity appeared dissociable from TMS-induced oscillatory activity, which tends to spread more remotely.

In addition to TMS-evoked activation studies, a large body of research has focused on oscillatory signatures arising from macro- and micro-scale neural recordings. Among these studies has arisen the concept of natural frequencies of human cortical modules, suggesting that distinct regions of the cortex may naturally oscillate at distinct frequencies (Niedermeyer, et al. 1999). Of great interest is an expansion of the natural frequencies concept, reporting evoked dominant oscillations in specific cortical regions, with posterior regions naturally resonating at lower frequencies (~10Hz, alpha) and anterior regions at higher frequencies (~40Hz, gamma) (Rosanova, et al. 2009; Ferrarelli, et al. 2012).

While the above results suggest state-dependent differences in cortical oscillations and neuronal connectivity, that may be examined using TMS-EEG, the difference in evoked oscillatory activity between resting and active task states has rarely been investigated (Johnson, et al. 2012), let alone how this effect might differ across different stimulation sites. To further explore this, in the present study we tested two groups of subjects (N=12 each) while simultaneously applying single-pulse TMS and recording resultant EEG responses. One group was tested while subjects were at rest, similar to the previous reports, while a second group was tested while actively engaged in a simple sensorimotor task. We repeated the methods and analysis regimes of the prior studies, wherein TMS was administered and frequency spectra data were obtained to measure the maximum evoked frequency at each stimulation site (Rosanova, et al. 2009).

## Materials and methods

### Subjects

Twenty-four right-handed subjects (12 females, age 19–36 years) with normal or corrected-to-normal visual acuity participated in this study, and were randomized into one of two groups: twelve subjects participated in the passive experiment (7 females, mean age 22.5 years), while the other twelve subjects participated in the active experiment (5 females, mean age 25.2 years). Following safety and ethical guidelines (Rossi, et al. 2009), all subjects were eligible to receive TMS. All subjects provided informed consent and all protocols were approved by the George Mason University Institutional Review Board.

### TMS

A focal figure-of-eight coil with 70mm wing diameter driven by a Magstim Rapid² biphasic stimulator (Magstim Inc., Wales, UK) was used to non-invasively stimulate the subjects’ cortex. On the right hemisphere of the scalp, three cortical sites were selected over the EEG electrodes P4 (occipital), C4 (parietal), and F4 (frontal). These sites were chosen as homologous regions to those stimulated previously in a study demonstrating distinct frequency responses (Rosanova, et al. 2009). To verify anatomical locations of the cortical stimulation sites a T1-weighted MRI of one subject was used and targeted using MNI coordinates in the Brainsight neuronavigation system (Rogue Research Inc., Montreal, Canada). MRI scans were unavailable for the other subjects in this study.

### High-density EEG recording during TMS

TMS-evoked potentials (TMS-EP) were recorded using an actiCHamp 64-channel amplifier (Brain Products GmbH, Germany) and TMS-compatible actiCAP slim active electrodes (international 10-20 system), with FCz as the online reference. BrainVision Recorder (v. 1.20.0801) was used to digitize the EEG at a sampling frequency of 5000 Hz. Electrode impedances were kept <20 kΩ. To minimize contamination of auditory potentials evoked by the click associated with the TMS coil discharge (ter Braack, et al. 2015), subjects underwent a TMS-click auditory perception (TMS-CAP) test prior to beginning the experiment. During the test, subjects wore inserted wired silicone-tipped earplug headphones with a Noise Reduction Rating (NRR) of 26 dB, while a masking noise with the same spectral profile of the TMS coil click was continuously played. Recordings of the TMS coil click emitted by the Magstim coil were used to create the masking noise and scrambled into a continuous sound file with the same spectral properties, thus, capturing the specific time-varying frequency components of the TMS click. For the TMS-CAP test, subjects listened to the masking noise while a brief TMS burst was administered on top of the FCz (average reference) electrode. Subjects were instructed to notify the experimenter if they could hear the TMS coil click. If a subject reported hearing the click, the volume of the masking noise was raised to a level still comfortable for the subject and/or the stimulator intensity output percentage was lowered until the click was as imperceptible as possible without lowering the stimulator output to an ineffective intensity (<40 V/m) (Rosanova, et al. 2009). Once the TMS-CAP was complete, subjects were required to continue wearing the earplug headphones for the duration of the experiment while listening to the masking noise at their individualized fixed volume. All TMS-induced artifacts were attended to during offline analysis.

### TMS protocol

During this experiment, subjects in both groups received a single-pulse TMS protocol. This consisted of a series of TMS pulses that were administered one-at-a-time over a part of the brain on the right hemisphere of the scalp, approximately over electrodes P4, C4, and F4 (Fig 1). At each of the three electrode sites, a total of 100 pulses were administered repetitively at each site, separated by a short period of time based on randomized experimental group assignment (passive or active), resulting in a total of 300 pulses overall. Based on their individual TMS-CAP results, subjects in the passive experimental group received single pulse stimulation at a fixed intensity of between 30–50% of maximum stimulator output (MSO; range 44.7–74.5 V/m), while subjects in the active experimental group received single pulse stimulation at a fixed intensity of between 35–50% of MSO (range 52.1–74.5 V/m,); both groups received an average of 42.5% MSO (± 7.23 for passive and ± 5.84 for active, respectively). According to an independent samples t-test, MSO values were not different between the two groups; *t*(22) = 0, *p* = 1.

**Fig 1.**
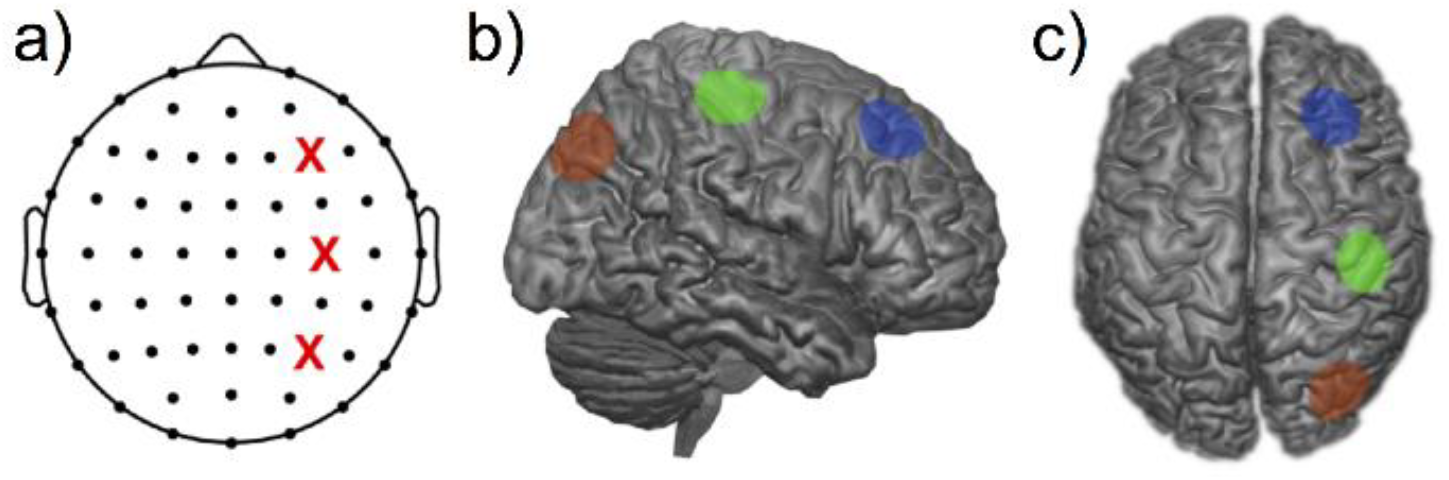
Stimulation sites. a) represents electrode stimulation sites, indicated by red X mark; top X is F4, middle X is C4, and bottom X is P4. Images b) and c) represent stimulation sites on a rendered brain as determined by Brainsight localization of the TMS coil on a sample subject.

We additionally calculated the modeled electrical field of our TMS stimulator across the different MSO intensities employed. To that end, we conducted electrical field modeling using SimNIBS software (v. 2.1.20) (Thielscher, et al. 2015), on a standardized MNI template with a modeled Magstim TMS coil matching our own. Similar to MSO values, we observed no differences between groups (Passive: average 63.3±10.77 V/m; Active: average 63.3±8.71 V/m). Additionally, we note that the lowest intensity in our tested sample (44.7 V/m) did not exceed the minimal intensity previously reported as minimal for evoking dominant frequencies (Rosanova, et al. 2009).

### General experimental procedures for both experimental groups

During the experiment, subjects sat in an ergonomic chair, relaxed, and with eyes open looking at a fixation cross on a screen. Once the selected electrode site was targeted, we stimulated it at an intensity that was set by the subject’s TMS-CAP results.

### Passive group experimental procedures

Subjects in the passive experimental group did not perform a sensorimotor task; instead, they were instructed to rest with their eyes open during the experiment. Subjects viewed an LCD monitor with a 120 Hz refresh rate (Cambridge Research Systems, United Kingdom) approximately 70-cm away with a black background, and were instructed to keep their eyes fixed on a 5×5 cm black fixation cross that appeared in the center of the screen. Each of the three electrode sites were stimulated in a counterbalanced block design and received 100 stimuli per block at a randomized inter-stimulation-interval (ISI) between 4–6 seconds.

### Active group experimental procedures and task

Subjects in the active experimental group performed a sensorimotor luminance detection task that relied on gradual signal detection, and was programmed in PsychoPy (v. 1.85.6) (Peirce, 2009). Subjects viewed a screen approximately 70-cm away with a grey background and a 5×5 cm fixation cross with a white outline in the center of the screen that gradually changed from a solid black center to solid white, achieving full luminance (Fig 2). Once the subject perceived the fixation cross to reach full luminance, they pressed a key with their right hand, upon which a single-pulse stimulation was delivered before the next trial automatically began. A white outline of a fixation cross was presented to each subject at the start of each trial, during which the interior gradually increased in luminance at a rate of 0.0025 value/frame (in HSV units). Each of the three electrode sites were stimulated in a counterbalanced block design and received 100 stimuli per block.

**Fig 2.**
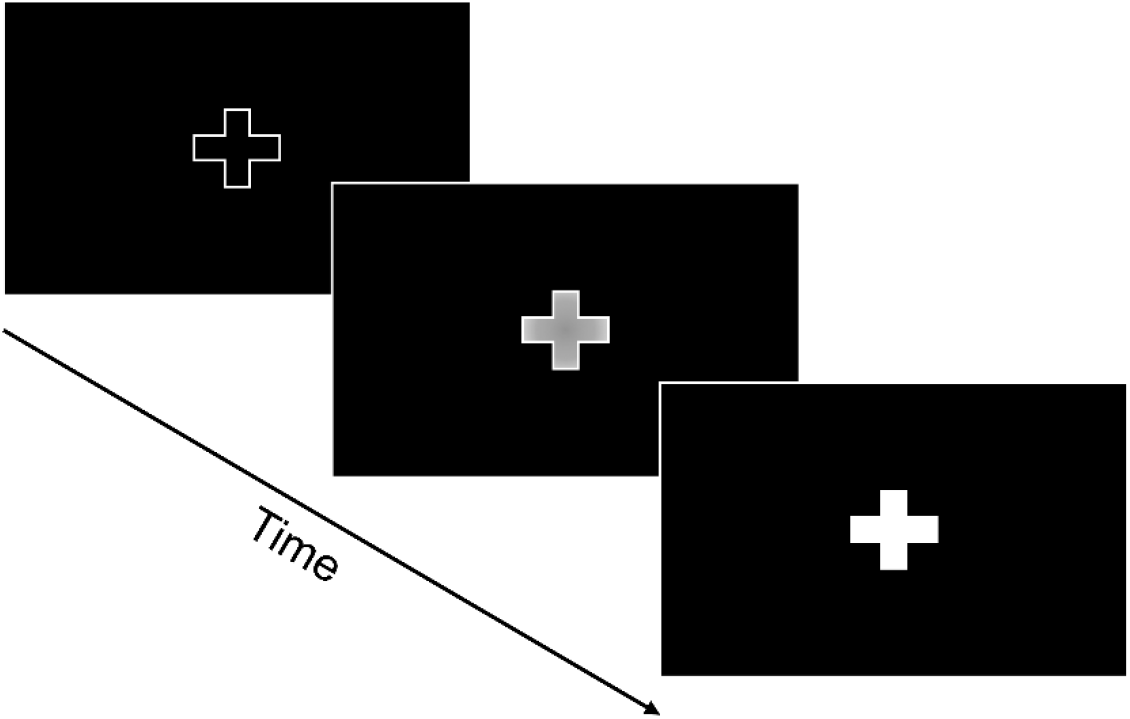
Schematic of the sensorimotor luminance detection task for the active experimental group. Subjects viewed a fixation point that gradually became illuminated in the center, and were required to press a button when they judged the luminance to be fully saturated, thus initiating the next trial.

### Analysis

Offline data analysis was conducted using the EEGLAB MATLAB Toolbox (Delorme & Makeig, 2004) in MATLAB R2017b (The MathWorks Inc., Natick, MA). Continuous data were downsampled to 500 Hz and subsequently analyzed via the *clean_rawdata* plugin (v. 0.34) (Kothe, 2014) to clean continuous data following the Artifact Subspace Reconstruction (ASR) method (Mullen, et al. 2013) to remove bad EEG channels. To prevent result biases from potentially removing excessive datapoints, *clean_rawdata* provided us with controlled, objective rejection criteria to eliminate noisy channels for such artifacts as eye blinks and face/neck muscle activity. Following this, all data were re-referenced to the grand average of all electrodes and then epoched for all three stimulation sites from −1000 to +1000 ms around the TMS pulse; the data for each epoch was baseline-corrected to the mean of the entire epoch span.

### TMS-artifact removal

Given the emergence of concurrent TMS and EEG as an important tool for assessing cortical properties, TESA—an open-source extension for EEGLAB—was created with the purpose of improving and standardizing analysis across the field of TMS-EEG research (Rogasch, et al. 2017). We applied the TESA toolbox to all three stimulation site epochs to remove artifacts; all steps adhered to the TESA pipeline (Rogasch, et al. 2017). This process involved 1) removing all data around the TMS pulse from −10 to +10 ms, 2) interpolating removed data within the TMS pulse, 3) removing noisy trials via EEGLAB’s built-in joint probability detection, 4) running a first round of independent component analysis (ICA) using the FastICA algorithm, 5) removing artifact components via visual inspection, 6) applying a first-order Butterworth filter with a bandpass of 1–100 Hz, as well as a notch filter to remove 60 Hz electrical line interference, 7) running a second round of FastICA with subsequent artifact component rejection. Following the above steps, data were again filtered between 1 and 50 Hz and segregated into separate, site-specific epochs.

### Time/frequency analysis

To analyze time-frequency domain responses we calculated the event-related spectral perturbation (ERSP) values based on Morlet wavelets, via the EEGLAB *newtimef* function, by convolving a mother wavelet at 100 linearly-spaced frequencies spanning 5 to 50 Hz, with 3.5 cycle wavelets and a 0.5 scaling factor. Baseline correction was applied to the average power across trials by subtracting the mean baseline power. Analysis of time/frequency data thus proceeded at the “global” level, following the convention of previous experiments (Rosanova, et al. 2009). Accordingly, global effects were determined by averaging, for each subject, the time/frequency spectrogram across all electrodes to form a single representation of the ERSP across the scalp. To minimize the effect of possible artifacts occurring at the time of stimulation, natural frequencies were calculated by averaging the ERSP values in a time window between 20 and 200 ms (see below).

### Global field power

In addition to the analysis of ERSP data, we also calculated global field power (GFP), defined as the reference-independent response strength, and calculated as the standard deviation across all electrodes at each timepoint (Murray, et al. 2008). GFP data were analyzed across all three sites of stimulation, separately for passive and active groups, in order to determine if there were any differences in evoked activity following TMS at any site.

### Natural frequencies

Our analysis of natural frequencies proceeded according to the description from previous reports (Rosanova, et al. 2009; Ferrarelli, et al. 2012). To determine the natural frequency for each subject at each stimulation site, the global ERSP (gERSP) response was analyzed by calculating the sum of power values for each frequency within the 20–200 ms time window, and then determining which frequency had the highest value. In this way, the max frequency would not be driven alone by a single frequency with a very high peak, but could instead be provided by a frequency with a moderate yet sustained response that was larger than at other frequency bands. Natural frequencies were calculated for each stimulation site for each subject in both groups.

### Statistical analysis

All statistical analysis of behavioral data and natural frequencies were carried out in SPSS (v. 19, IBM Corporation). For the analysis of global and local effects, we employed cluster-level corrections for significance (*p* < 0.05) (Maris & Oostenveld, 2007) and implemented via Fieldtrip using the *statcondfieldtrip* command in EEGLAB. For both local and global effects, we determined regions of significant deviation from baseline for each of the sites, for each of the two groups. In addition, we compared the gERSP between groups, by averaging across all three sites within each group and comparing the overall responses.

## Results

### Global response to TMS

Our initial analyses set out to attempt to reproduce the methods used by previous studies (Rosanova, et al. 2009; Ferrarelli, et al. 2012) for reporting dominant frequencies in specific cortical regions. In these studies, an increase in power is observed following TMS that is maximal at a particular frequency band, dependent on the site of stimulation. These changes reflect the spectral properties of the TMS-evoked oscillations, which consists of a number of repeated positive and negative deflections (Lioumis, et al. 2009). When examining the gERSP response, averaged across all electrodes, we observed a combination of increases and decreases in power following TMS. Notably, only the *decreases* in power survived our cluster-corrected significance threshold, in contrast to the original findings. This finding was observed across both passive and active groups. However, we note that our study used a different design and methods to these previous reports. Most notably, these previous studies stimulated cortical regions at a much higher intensity than ours; in the present study, we sought to reduce the impact of artifact peripheral components in the TMS-EP (Conde, et al. 2019). In doing so, our stimulation intensities were far lower than that used previously. Nevertheless, the difference in evoked frequency response between conditions, our modeling findings, and behavioral differences in the active group between motor cortex stimulation and other sites suggest that our stimulation intensities were sufficient for inducing activity in cortical columns.

Crucial differences were observed between passive and active groups, as well as between the different sites of stimulation. For the passive group, we observed decreases in power that were synchronous with the TMS pulse in the gamma frequency band (40–50 Hz) across all three sites. Across stimulation sites, the gamma desynchronization became longer lasting from posterior to frontal regions, and was further accompanied at the frontal site by a significant decrease in the high beta range (20–30 Hz) approximately 100–300 ms after the TMS pulse. In contrast, the active group exhibited a larger desynchronization response across all three sites, extending from the beta to gamma range (Fig 3).

**Fig 3.**
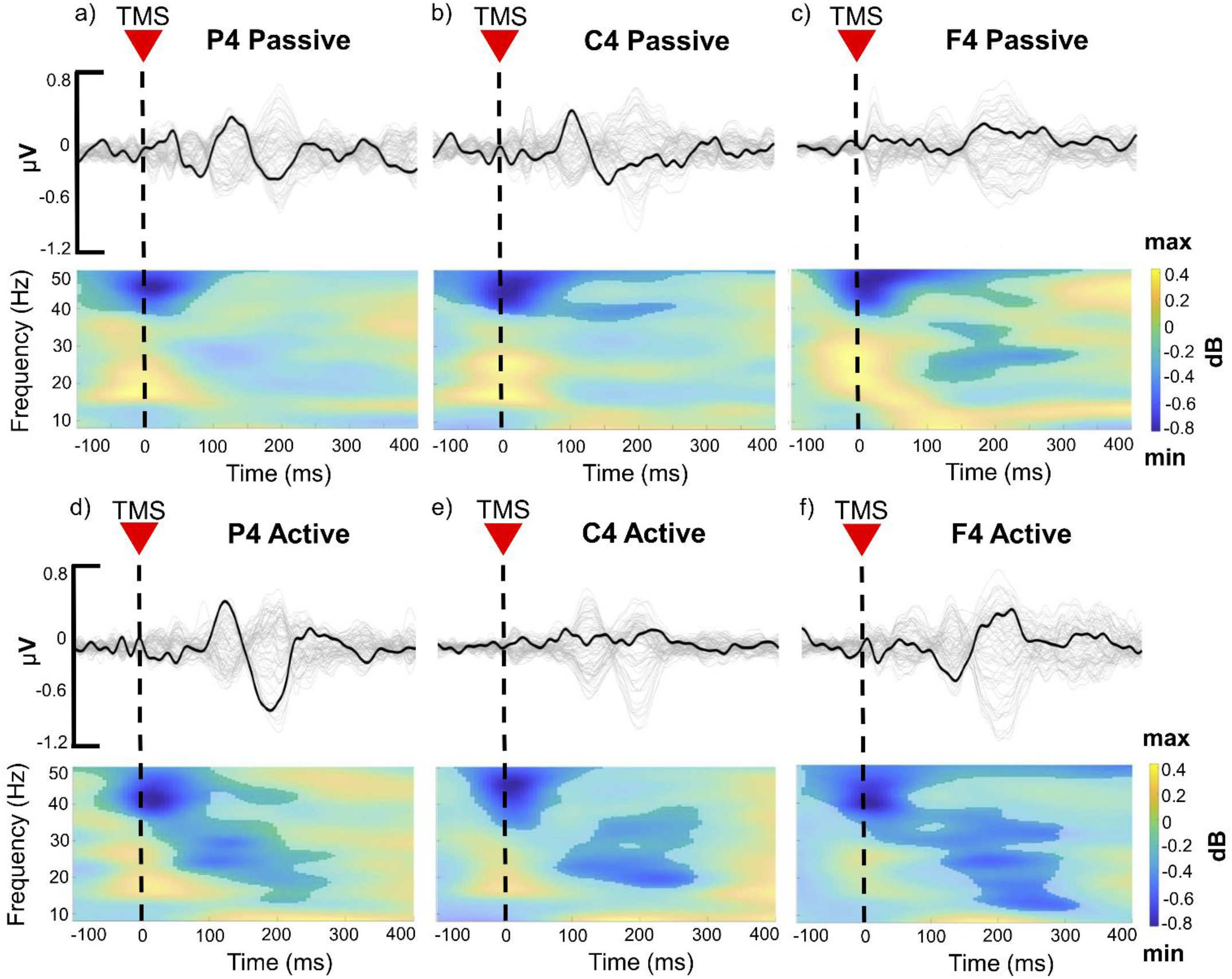
Global plots for all subjects illustrating the three cortical sites targeted by TMS. Butterfly plots (top panels) of all electrode time courses with the black trace line highlighting the electrode directly underlying the stimulator. ERSP plots (bottom panels) display saturated color areas representing significant frequency (Hz) activation compared to baseline. (a) P4 for passive group, (b) C4 for passive group, (c) F4 for passive group. (d) P4 for active group, (e) C4 for active group, (f) F4 for active group. Significance was determined via cluster-based permutation testing.

In addition to the spectral response, we also calculated and measured the GFP. Here, as well, we observed no differences between site in the evoked response, nor was there any difference between passive and active groups (Fig 4).

**Fig 4.**
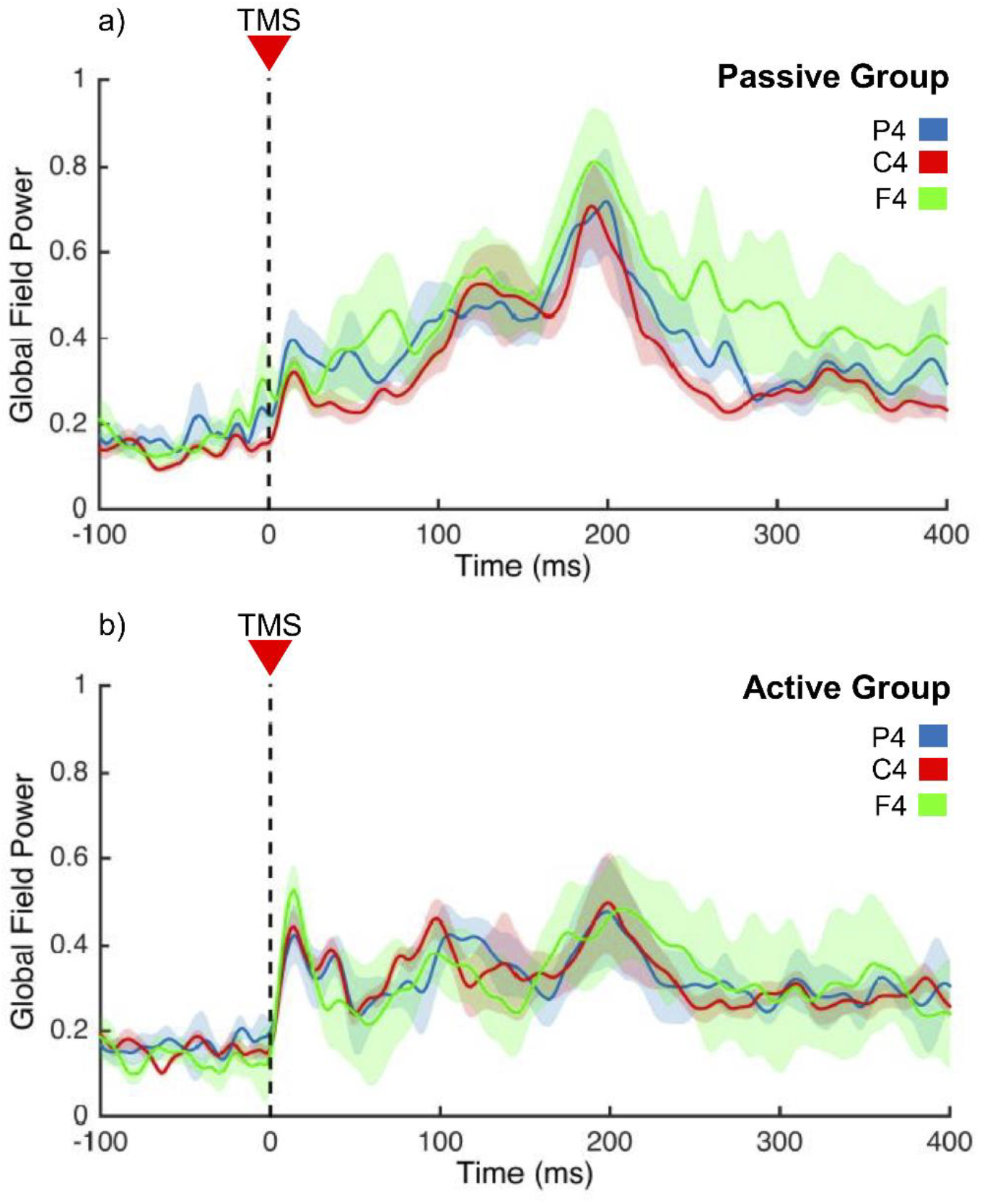
Global field power (GFP) across stimulation sites and groups. Shaded regions display standard error. No differences between stimulation site or group were detected.

### Natural frequencies

The major finding of the previous work was that the so-called “natural frequency,” characterized as the frequency band with the largest sustained response to TMS, increases in a rostro-caudal gradient. Calculating the natural frequency using the same method outlined by the previous authors yielded a range of values across all three sites. Though the individual maximum frequencies showed a large oscillatory range, there were no outliers. Yet, no linear effects were observed in these values across all three sites, for either the passive or active groups. However, along with the gERSP responses, we observed that the active task group exhibited significantly higher natural frequencies evoked by TMS than the passive group. A two-way ANOVA revealed that group type had a significant effect on mean activation in Hz (*F*(1, 22) = 4.557, *p* = 0.044, ηp^2^ = 0.172), with higher frequencies reported during the active experimental group (*M* = 22.43, *SD* = 12.24) compared to the passive experimental group (*M* = 15.66, *SD* = 12.6).

### Active group response times

Response times (RTs) were calculated in seconds (s) for the active group, which performed the illumination detection task; P4 (2.20±1.24), C4 (2.44±1.27), F4 (2.26±1.26) (Fig 6). A nonparametric Friedman test showed a significant difference in RTs between stimulation sites; *χ*^2^(2, 12) = 8.667, *p* = 0.013. Post hoc analysis with Wilcoxon signed-rank tests were conducted with a Bonferroni correction applied, resulting in a significance level set at *p* < 0.05. Median (IQR) RTs based on stimulation sites were 2.13 s (1.19 to 2.74) for P4, 2.05 s (1.52 to 2.92) for C4, and 2.06 s (1.17 to 2.76) for F4, respectively. There were no significant differences between RTs for P4 and F4 (*Z* = −0.784, *p* = 0.433) or between RTs for C4 and F4 (*Z* = −1.412, *p* = 0.158). However, there was a significant difference between RTs for C4 compared to P4, with slower RTs overall for C4 (*Z* = −1.961, *p* = 0.05).

**Fig 5.**
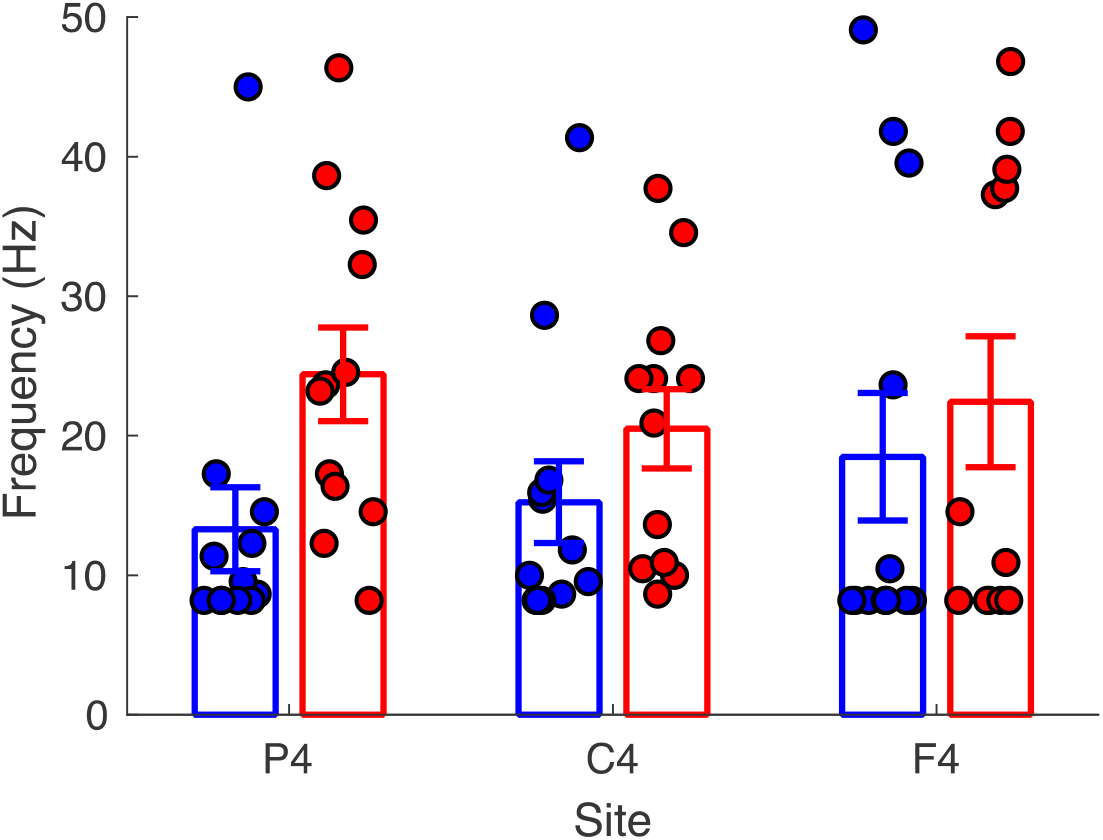
Mean natural frequencies, based on stimulation site and group assignment. Individual data points represent the maximal evoked frequency for each subject in the resting (blue) and active (red) state subject groups. Consistently higher evoked frequencies were observed across all stimulation sites for the active state subjects. Error bars represent standard error.

**Fig 6.**
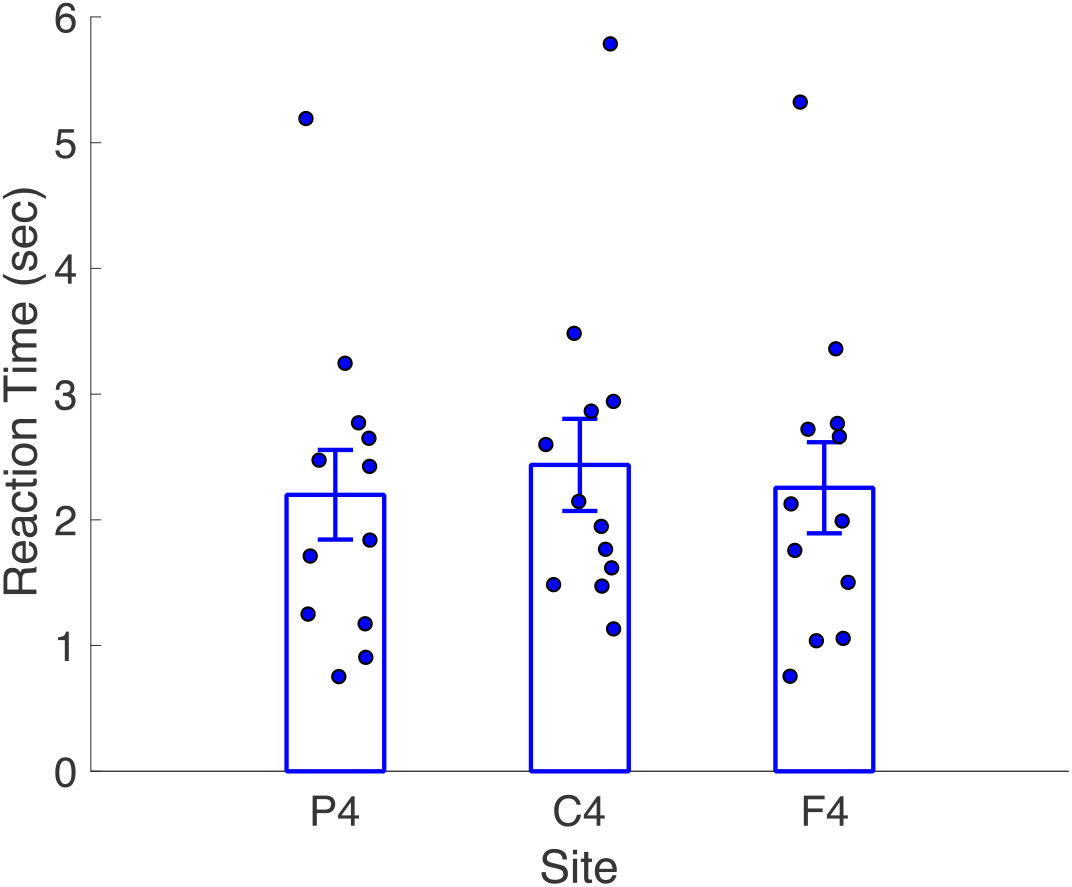
Average active group task response times (RTs) in seconds, based on electrode stimulation site. Blue circles represent for mean RTs for each active task subject, per stimulation site. Subjects were moderately slower in responding following single-pulse stimulation over C4. Error bars represent standard error.

## Discussion

In the current study, we used concurrent TMS-EEG to investigate cortical reactivity differences between passive state and active sensorimotor task performance. Our results revealed a complexity of patterns in global and local changes that occur in response to stimulation, in addition to conspicuously broader and distinct patterns between the passive and active states. These findings suggest cortical regions exhibit complex frequency-specific profiles. Specifically, our findings suggest that oscillatory mechanisms are characterized by more complex, state-dependent patterns than have previously been appreciated. Additionally, our findings suggest that patterns of ongoing spontaneous activity are modified by task performance, and differ based on individual activation patterns.

Endeavoring to explain the complexity of varying oscillatory bands, further investigation into whether timescales of different frequency bands correlate with a sensory-to-higher processing hierarchy was conducted (Mellem, et al. 2017). However, no strong biases toward specific timescales across cortical regions was observed, nor exclusivity of lower areas biased toward faster frequencies or higher areas biased toward slower frequencies. Thus, it was contended that dominant higher-frequency bands observed in the frontal cortex during concurrent TMS-EEG studies may be a result of that brain region’s involvement in higher level cognition. This explanation has potential to elucidate our findings of higher frequency evoked power across the brain during sensorimotor task engagement. Furthermore, frequency bands may serve as channels of communication across brain regions, though dependent on the activation in multiple bands within each region (Hillebrand, et al. 2016).

Perhaps the most notable aspect of our findings is the difference in evoked frequency between the experimental groups. While performing the sensorimotor task, evoked responses became more widespread in both frequency and time; additionally, when comparing the natural frequency between experimental groups, the sensorimotor task was observed to evoke a consistently higher frequency than in the resting state group. This difference suggests that cognitive engagement incorporates higher frequency oscillations, consistent with several other known findings of brain function (Crone, et al. 1998; 2011; Canolty, et al. 2006; Voytek, et al. 2010; Groppe, et al. 2013). A state-dependent TMS effect might account for these findings. State-dependency is defined as response changes according to the state of the cortex when the stimulus, such as a TMS pulse, is applied (Siebner, et al. 2009). Moreover, the state of activation has been shown to influence the response (Romero, et al, 2019;Silvanto, et al. 2007;2008; Romei, et al. 2008; Massimini, et al. 2010;Miniussi, et al. 2010; Romei, et al. 2016; Petrichella, et al. 2017). The effect of small TMS pulses might be facilitated if the cortex is already active; thus, it would be reasonable to presume that single pulse stimulation may well enhance cortical activation while a subject is actively engaged in a task (Matthews, 1999). Previous studies have shown evidence of the effect of TMS pulses varying as a function of the state of the brain. For example, when comparing neuronal activation during resting/baseline states to active task engagement, researchers found TMS over the motor cortex enhanced activation during motor execution and motor imagery (Kasai, et al. 1997; Fadiga, et al. 1999; Hashimoto, et al. 1998), while others found greater ease of inducing phosphenes with TMS over the occipital region during visual mental imagery (Sparing, et al. 2002). The latter finding suggests distinct operational modes for the brain between resting state and task-based networks. Consistent with this view, previous investigations comparing resting-state and task-based network activity in functional magnetic resonance imaging (fMRI) have revealed network reorganization between these states (Spadone, et al. 2015; Gonzalez-Casillo & Bandettini, 2018); in particular, the frequency profile of fMRI inter-connectivity shifts between resting and task-based activity, with lower frequencies dominating the former and more broadband representation during tasks (Ciuciu, et al. 2014). Although these fluctuations operate on an order of magnitude below those measured by EEG in the present study (0.01-0.1Hz), they reveal a similar pattern to our findings, suggesting a correspondence (Bridwell, et al. 2013).

Notably, for the current study, the higher evoked frequencies did not depend on the stimulation site, suggesting a global change in brain functioning, independent of the local changes. Finally, our findings confirm that TMS can be a useful tool for evoking latent oscillations in the brain [14].

### Limitations

In the current study, there is a possible limitation that should be noted. We recognize that the stimulation levels in our study are lower, on average, than used in previous reports. This was done to avoid auditory and somatosensory evoked artifacts in the EEG response. As a result, the evoked responses observed in EEG spectra are lower than previously reported. Yet, we note that TMS intensity was above 40 V/m, previously reported as minimal for evoking dominant frequencies (Rosanova, et al. 2009). Further, while the evoked responses are lower, they still adhere to the overall shape of the TMS-EP, exhibiting a clear N1-P2 complex. Additionally, if stimulation was having no effect, then no difference should be expected 1) between groups, or 2) for C4 stimulation on RT in the active-state, both of which were observed.

As an additional note, in observing our findings, one may discern that the frequency spectra for the global response exhibits only significant decreases in power, with a greater spread for the active group, whereas an overall higher natural frequency was observed for the active group. This apparent discrepancy can be explained by differences in the analyses; in the global frequency analysis (Figure 3), significance is assessed against baseline, in which only decreases were found to exceed the threshold. In contrast, the natural frequency analysis does not look for significant differences versus baseline, but only considers which frequency band showed the biggest consistent increase. As such, both analyses approach the present data with different outcomes in mind.

Finally, we note the marginal difference in stimulation rate between active and passive groups. In the passive group, stimulation was repeated at a randomized rate of 0.16 – 0.25Hz, whereas in the active group, because stimulation was tied directly to subject responses, a rate of 0.4 – 0.45Hz occurred. While the active subjects received a higher rate of stimulation, it is unlikely to have contributed to the observed differences between active and passive groups. First, so-called “slow” rTMS (<0.5 Hz) has only demonstrated inconsistent effects (Hoffman & Cavus, 2002), and with differences observed between lower or higher ranges. Second, while slower rates of rTMS can affect cortical responses, these are only administered in a steady, rhythmic fashion, and for a far higher number of pulses than used here per stimulation site. Third, in a post-hoc analysis, we found that faster RTs (and thus faster stimulation rates) did not correlate between subjects with differences in the observed natural frequency^*^. Lastly, within subjects, we note that C4 stimulation, which led to a slower RT and thus slower stimulation rate, similarly did not engender lower natural frequencies.

## Conclusions

We investigated TMS-evoked cortical reactivity differences between subjects who were either at rest (passive group) or engaged in a sensorimotor task (active group), while recording resultant EEG responses. The differences in evoked responses between the two experimental groups suggests that oscillatory mechanisms are characterized by complex, state-dependent patterns, with an overall higher mode of frequency during active engagement.

## Acknowledgements

We would like to acknowledge Melody Barnard, our former undergraduate research assistant, for her help during data collection.

## Footnotes

* Pearson correlation: r = −0.051, p>0.05

